# Complexity measures of the electroencephalograph capture loss and recovery of consciousness in patients anesthetized with propofol

**DOI:** 10.1101/594002

**Authors:** Sarah L. Eagleman, Divya Chander, Christina Reynolds, Nicholas T. Ouellette, M. Bruce MacIver

**Author notes:** These authors contributed equally to this work (Sarah L. Eagleman, Divya Chander).

## Abstract

Propofol is one of the most widely used anesthetics for routine surgical anesthesia. Propofol administration alone produces EEG spectral characteristics similar to most hypnotics; however, inter-individual variation can make spectral measures inconsistent. Complexity measures of EEG signals could offer universal measures to better capture anesthetic depth as brain activity exhibits nonlinear behavior at several scales. We tested the potential of nonlinear dynamics analyses to identify loss and recovery of consciousness at clinically relevant timepoints. Patients undergoing propofol general anesthesia for various surgical procedures were identified as having changes in states of consciousness by the loss and recovery of response to verbal stimuli after induction and upon cessation of anesthesia, respectively. Nonlinear dynamics analyses showed more significant differences between consciousness states than most spectral measures. Thus, complexity measures could provide a means for reliably capturing depth of consciousness based on subtle EEG changes at the beginning and end of anesthesia administration.

## INTRODUCTION

Since the 1990s, commercial electroencephalogram (EEG) devices have been introduced into the operating room to monitor depth of anesthesia in surgical patients. Research has demonstrated that the use of these devices reduces the risk of intraoperative recall in patients (Avidan et al., 2013; G. A. Mashour & Avidan, 2015; Myles, Leslie, McNeil, Forbes, & Chan, 2004; O’Connor et al., 2001). Additionally, monitoring anesthetic depth is becoming increasingly important, since lighter planes of anesthesia speed recovery times, decrease operative costs (Dexter, Macario, Manberg, & Lubarsky, 1999) and reduce morbidity, including the potential for post-operative delirium and cognitive dysfunction in vulnerable populations of surgical patients (B.A. Fritz, Maybrier, & Avidan, 2018; Bradley A. Fritz et al., 2016; Kertai et al., 2011; Petsiti et al., 2015; Tarnal, Vlisides, & Mashour, 2015); though see also (Kalkman, Peelen, Moons, Group, & Group, 2011; Wildes et al., 2019) for limitations. However, lighter planes of anesthesia impart a greater risk of intraoperative recall in patients (Avidan et al., 2013; G. A. Mashour & Avidan, 2015). Commercially available monitors have consistently been shown to lack the sensitivity, accuracy, and response kinetics needed to prevent intraoperative awareness (Avidan et al., 2013; Health Quality Ontario, 2004; Schneider, Gelb, Schmeller, Tschakert, & Kochs, 2003), and different measures from different monitors produce significantly different processed EEG assessments at equal effect site concentrations (Soehle et al., 2008). Improvements to brain monitoring may be sought in new analytical techniques that capture the full nonlinear structure of the EEG signal instead of the traditional spectrally-derived metrics.

EEG signals derived from the frontal cortex show stereotypic responses to GABAergic anesthetics (e.g. propofol and sevoflurane), and frequency domain measures are particularly sensitive to anesthetic levels (Billard, Gambus, Chamoun, Stanski, & Shafer, 1997; Ching, Cimenser, Purdon, Brown, & Kopell, 2010; Patrick L Purdon, Sampson, Pavone, & Brown, 2015; Rampil, 2001; Vijayan, Ching, Purdon, Brown, & Kopell, 2013; Winters, 1976). For most anesthetics, loss of response (in humans, equated with loss of consciousness) occurs when low amplitude, higher frequency EEG waveforms are replaced with higher amplitude slow-wave patterns, similar to the delta rhythms seen during slow wave sleep (Chander, García, MacColl, Illing, & Sleigh, 2014; Patrick L Purdon et al., 2015). These delta rhythms are gradually replaced by burst suppression patterns as patients transition to deeper surgical planes of anesthesia (Pilge, Jordan, Kreuzer, Kochs, & Schneider, 2014). Interestingly, recovery of consciousness is associated with much less delta activity and higher amounts of low amplitude, fast activity on awakening, indicating a hysteresis of brain states for loss and recovery from anesthesia (Chander et al., 2014; Stanski, Hudson, Homer, Saidman, & Meathe, 1984), also termed ‘neural inertia’ which may reflect differential activation of neural networks during these states (Friedman et al., 2010; Tarnal et al., 2015). Emergence from general anesthesia also does not typically follow a stereotyped pattern, but rather a more heterogenous spectral pattern during the establishment of conscious awareness and responsivity (Chander et al., 2014). Further, frequency domain measures can differ for various classes of anesthetics and adjuvant agents (GABA versus non-GABAergic), in response to surgical stimulation, across age groups, and between people, making spectral domain measurements non-invariant.

The brain itself performs computations in both linear and nonlinear fashions, and both cognitive and arousal states have been characterized using analyses based on nonlinear dynamics (Ma, Shi, Peng, & Yang, 2018; Stam, 2005; Walling & Hicks, 2006; Watt & Hameroff, 1988; Widman, Schreiber, Rehberg, Hoeft, Elger, et al., 2000). An example of such a method is the use of time-delayed embeddings derived from EEG signals. Anesthesiologists have noted that such embeddings could distinguish anesthetic depth (Watt & Hameroff, 1988; Widman, Schreiber, Rehberg, Hoeft, & Elger, 2000) as well as recovery of consciousness in surgical patients (Walling & Hicks, 2006). Sometimes, time-delayed embedding signals are observed to explore only a limited region of all possible states in a multidimensional space in which they are embedded; instead, they settle onto attractors in state space, and can thus be seen as probes of the underlying dynamics. We have previously shown that attractors generated from microEEG (local field potential) signals in rodents are sensitive to isoflurane-induced loss of righting responses in rats (MacIver & Bland, 2014). Loss of righting reflex is a surrogate measure for loss of consciousness in rodents (Frank & Jhamandas, 1970). Specifically, we observed that initially spherical attractors became flattened and more ellipsoidal when moving from waking to loss of righting reflex (MacIver & Bland, 2014). We observed a similar change in attractor shape when characterizing human subjects anesthetized with a combination of remifentanil and nitrous oxide (Eagleman, Drover, Drover, Ouellette, & MacIver, 2018), and another patient population anesthetized with propofol and fentanyl (Eagleman, Vaughn, et al., 2018). We quantified the shape changes we observed using a novel geometric phase-space analysis, termed the ellipse radius ratio (ERR). We fit the 3D attractor with an ellipsoidal solid of revolution (Khachiyan, 1980) and then calculated the ratio between the major and minor axes. This ellipse radius ratio changes significantly and consistently before and after loss and recovery of response during several anesthetic regimes (Eagleman, Vaughn, et al., 2018; Eagleman, Drover, et al., 2018). Additionally, the ERR also shows a significant difference in adult patients across a wide range of ages (Eagleman, Vaughn, et al., 2018; Eagleman, Drover, et al., 2018). To make sense of this shape change, we compared this analysis to other complexity measures such as the correlation dimension and multiscale entropy, and found a significant correlation with multiscale entropy (Eagleman, Vaughn, et al., 2018). Characterization of attractors using correlation dimension (CD) has been proposed to provide a measure of complexity or information content of EEG signals (Walling & Hicks, 2006; Watt & Hameroff, 1988). Currently, it is unknown whether either correlation dimension or our geometric phase-space analysis is sensitive to recovery of consciousness for propofol anesthesia. It is also unknown how loss and recovery of response properties of EEG signals differ, and how the dynamics change with anesthetic administration and time as patients lose and regain their ability to respond (lose and regain consciousness).

To determine whether nonlinear dynamics analyses capture depth of consciousness reliably and to compare the dynamics of complexity changes with loss and recovery of response, we explore the ability of correlation dimension and our ellipse radius ratio (ERR) to capture the loss and recovery of response during propofol anesthesia. We compare our analyses from nonlinear dynamics to frequency-derived measures of EEG signals for loss and recovery of consciousness, and characterize the dynamics that occur during these transitions.

## RESULTS

### Spectral differences between pre- and post-LOR and ROR transitions are best captured by spectral edge frequency and gamma

We measured the EEG spectrum collected from F7 referenced to AFz in 20 s artifact-free clips identified pre- and post-loss (LOR) and recovery (ROR) of response (Figure 1A). Visualization of the EEG spectrum before and after loss of response (LOR) shows similar changes to what has been previously reported: notably, increases in lower frequencies and decreases in higher frequencies, including a visually apparent increase in the alpha range (8-12 Hz) (Breshears et al., 2010; Chander et al., 2014; P. L. Purdon et al., 2013) (Figure 2). In contrast, emergence from propofol anesthesia is marked by a more uniform distribution of power across frequency bands (Figure 2), which has also been reported previously (Chander et al., 2014). When we look at the spectral changes through time we observe the slower dynamic changes in the spectral content throughout LOR (Figure 1B, Figure 3). It appears that frequencies below 12 Hz gradually increase, while frequencies above 12 Hz gradually decrease and the dominant frequency, near the alpha (8-12 Hz) range, emerges near to and following the LOR transition (Figure 1B, Figure 3). However, this alpha increase does not reach statistical significance in this sample (Table 1). In this cohort of patients, the ROR dynamics appear more abrupt, with sharp changes in frequency content near the ROR transition (Figure 3). The more uniform power distribution across frequency bands occurs within seconds of the marked ROR event, and the previously dominant alpha rhythm is replaced by a more pronounced beta rhythm (Figure 3), though it does not reach statistical significance in this sample (Table 2).

**Figure 1.**
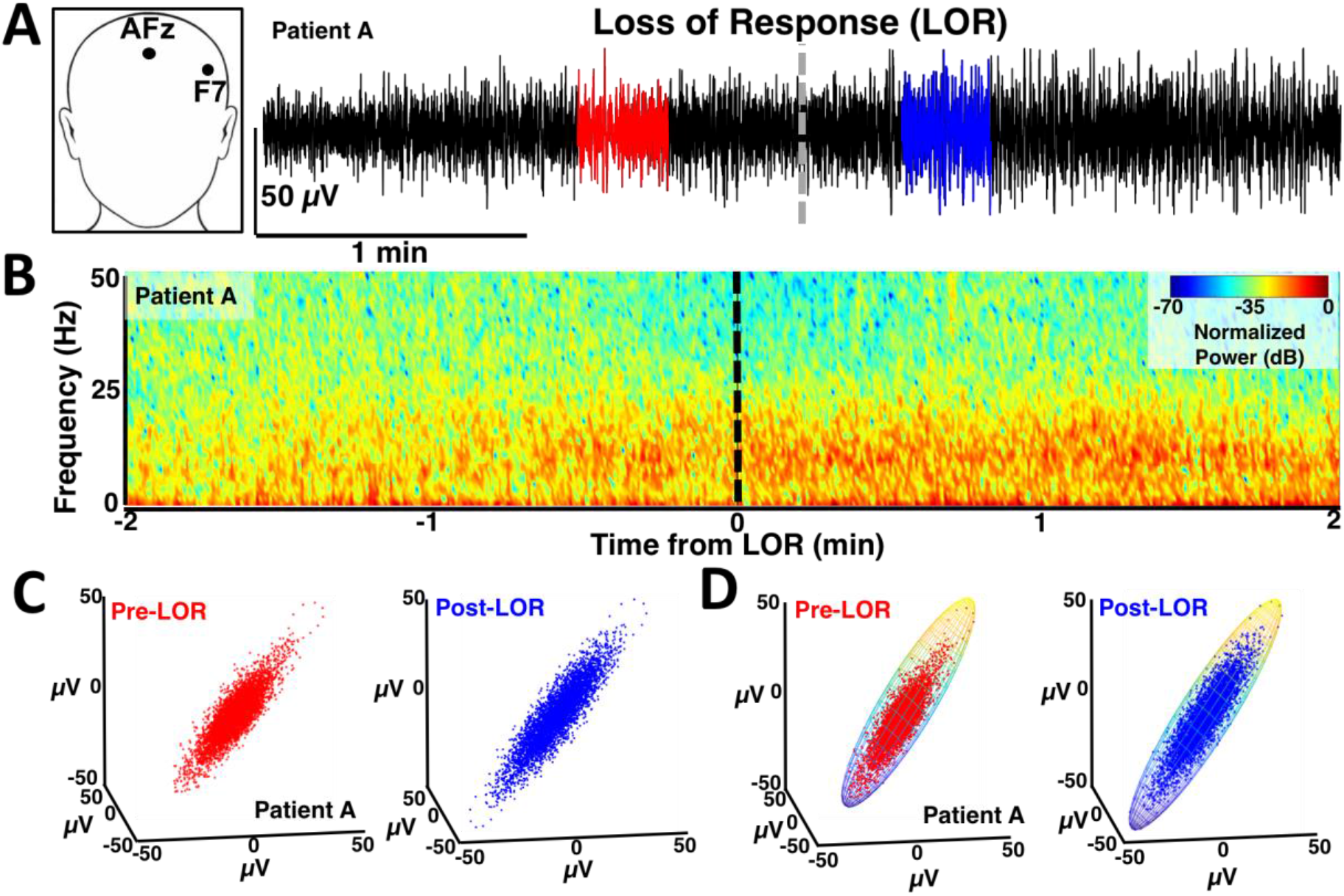
Example of activity around loss of response (LOR). **A)** EEG was recorded from F7 referenced to AFz. Twenty second artifact-free segments were identified before (red) and after (blue) loss of response (LOR) and used for subsequent analysis. Data shown is from patient A. **B)** Spectral activity around LOR shows increases in low frequency activity (< 12 Hz), previously termed slow-wave anesthesia, and decreases in high frequency activity (>12 Hz). **C)** Time-delayed embeddings generated from EEG activity from before (red) and after (blue) LOR. **D)** One way differences in the time-delayed embeddings pre- and post-LOR were quantified by fitting them with ellipsoids.

**Figure 2.**
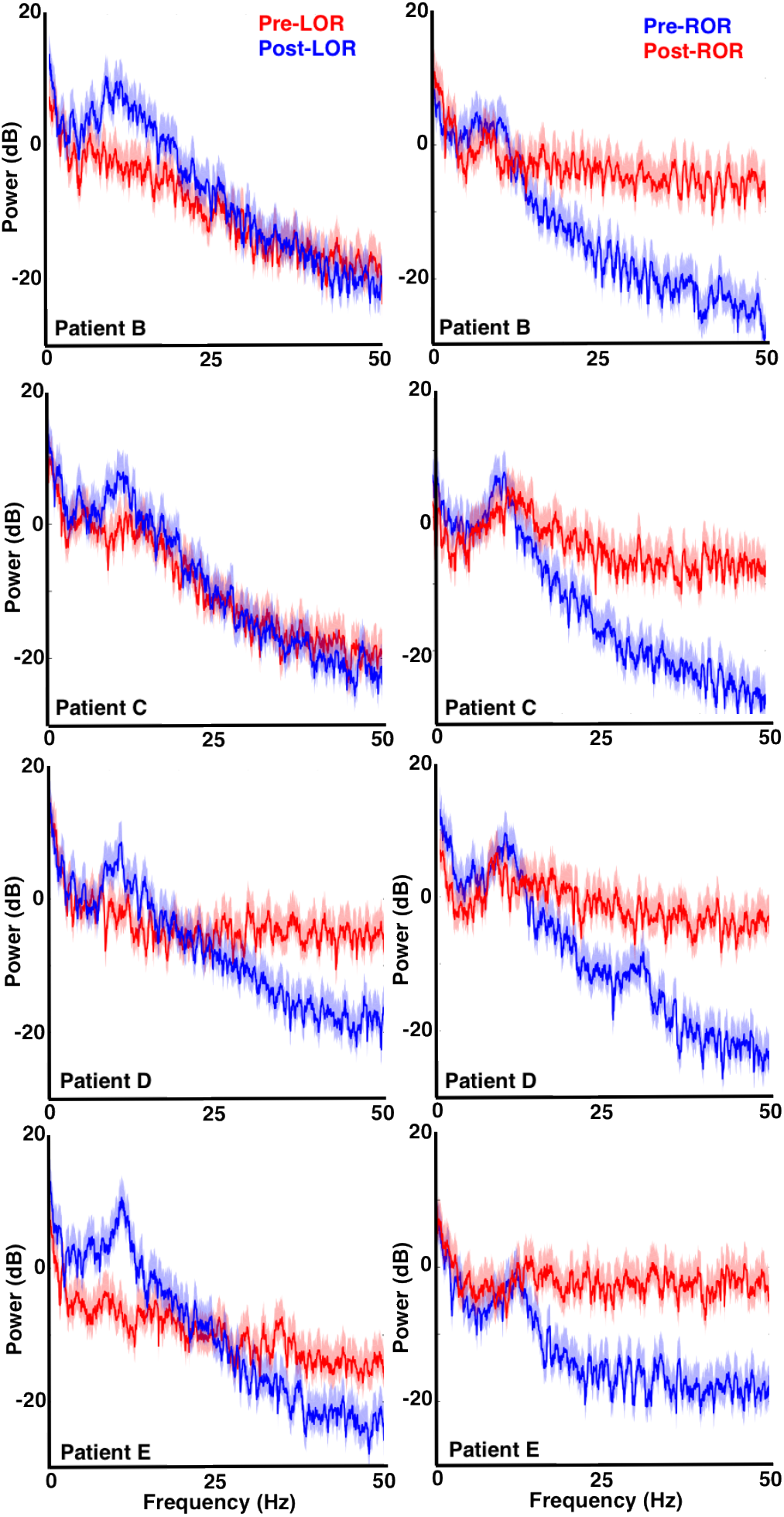
Power Spectra before and after loss and recovery of response. Power spectra around loss (left) and recovery (right) of response from four individual patients is shown. Awake or responsive periods (pre-LOR in left column and post-ROR in right column) are in red while anesthetized periods (post-LOR in left column and pre-ROR in right column) are in blue. The responsive state (red) is characterized by higher power in the frequency bands > 12 Hz, while the anesthetized or LOR state is characterized by higher power in the slower frequency bands < 12 Hz, with peaks in the delta (1-4 Hz) and alpha (8-12 Hz) band range, previously defined as slow-wave anesthesia (SWA). Solid red and blue lines indicate the mean value at each frequency, while the shading indicates a 95% confidence interval using a theoretical error distribution.

**Figure 3.**
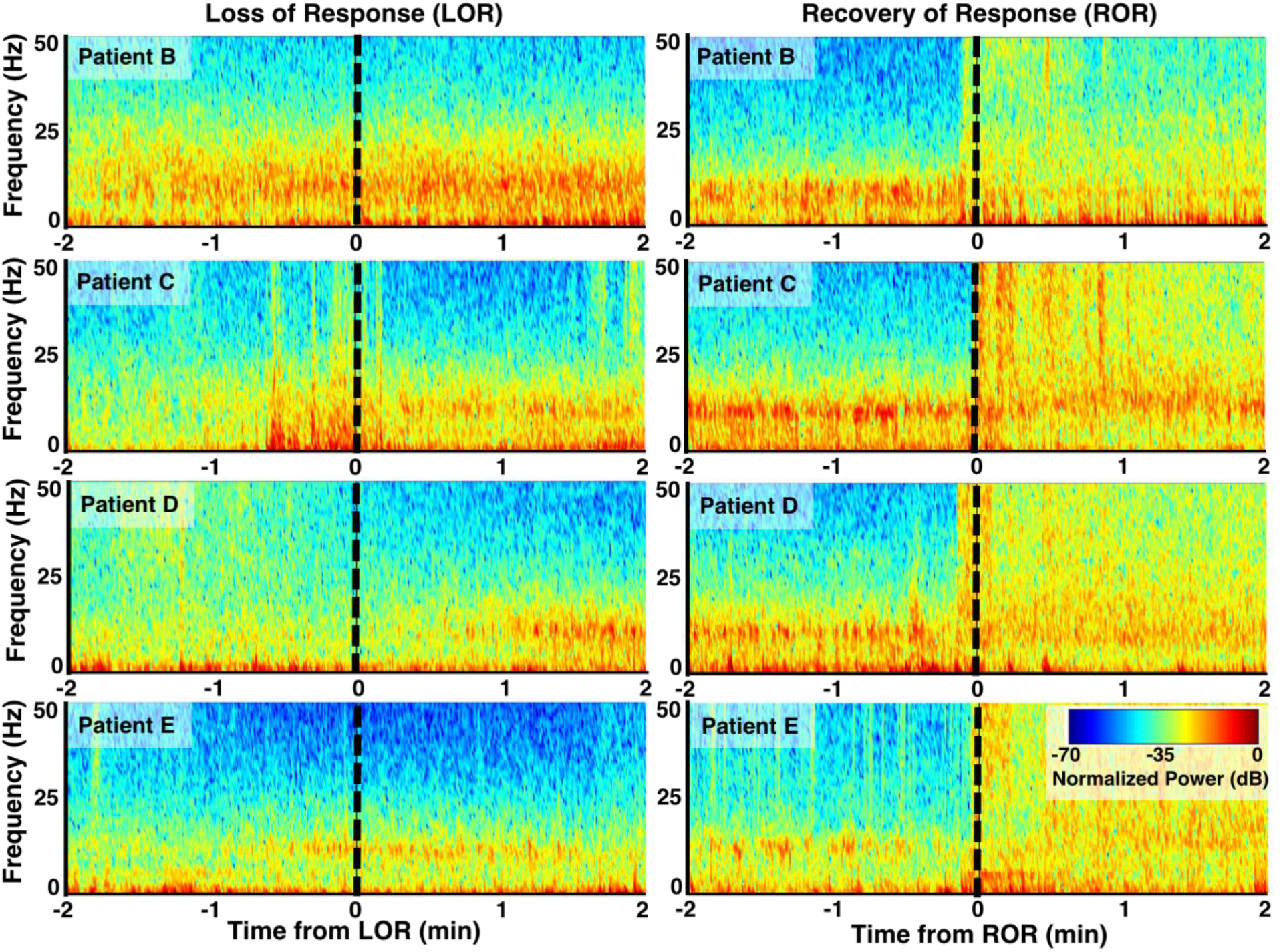
Spectrograms before and after loss and recovery of response. Examples of normalized spectrograms around loss (left) and recovery (right) of response events from four patients. Preceding and following LOR, the pattern of spectral activity is characterized by an increase in power of slow-wave activity, termed slow-wave anesthesia^18^, in both the delta (1-4 Hz) and alpha (8-12 Hz) bands; a predominant alpha rhythm is particularly noticeable. This transition emerges slowly during the LOR period, and with a variable time-course for each patient. In contrast, the period of emergence from general anesthesia to ROR is marked by a more uniform distribution of power across frequency bands, and the previously dominant alpha rhythm is replaced by a more pronounced beta rhythm. In the 4 patients shown, this transition appears to be more abrupt than the trajectory toward LOR.

**Table 1.** Summary of spectral changes from before and after loss of response (LOR). **Percentage** of power in individual frequency bands are reported as medians [25, 75 percentiles]. Significance values reported are from Wilcoxon signed rank tests, corrected for multiple comparisons. The * indicates that the percentage of power change was significant at *p* < 0.05 corrected.

| Frequency band | Percentage of Power<br>Pre-LOR | Percentage of Power<br>Post-LOR | Significance<br>value,<br>corrected |
| --- | --- | --- | --- |
| **Delta (1 – 4 Hz)** | 37.8% [31.2, 44.7] | 33.1% [28.7, 41.9] | 0.63 |
| **Theta (4 – 8 Hz)** | 10.8% [6.2, 16.2] | 9.6% [8.1, 12.2] | 0.63 |
| **Alpha (8 – 12 Hz)** | 9.4% [6.1, 11.9] | 15.6% [10.2, 19.9] | 0.075 |
| **Beta (12 – 25 Hz)** | 11.4% [8.1, 20.4] | 17.4% [9.3, 20.7] | 0.63 |
| **Gamma (25 – 50 Hz)** | 6.5% [4.0, 9.6] | 1.9% [1.2, 2.9] | 0.0006* |

**Table 2.** Summary of spectral changes from before and after recovery of response (ROR). **Percentage** of power in individual frequency bands are reported as medians [25, 75 percentiles]. Significance values reported are from Wilcoxon signed rank tests, corrected for multiple comparisons. The * indicates that the percentage of power change was significant at *p* < 0.05 corrected.

| Frequency band | Percentage of Power<br>Pre-ROR | Percentage of Power<br>Post-ROR | Significance<br>value,<br>corrected |
| --- | --- | --- | --- |
| **Delta (1 – 4 Hz)** | 44.5% [27.6, 64.6] | 26.2% [16.3, 44.7] | 0.010* |
| **Theta (4 – 8 Hz)** | 10.8% [9.6, 13.3] | 7.3% [5.6, 9.8] | 0.014* |
| **Alpha (8 – 12 Hz)** | 11.9% [6.1, 17.7] | 7.1% [4.9, 9.5] | 0.010* |
| **Beta (12 – 25 Hz)** | 10.6% [3.8, 18.4] | 17.8% [10.4, 28.8] | 0.049* |
| **Gamma (25 – 50 Hz)** | 1.3% [0.7, 4.3] | 11.8% [7.8, 22.4] | 0.0055* |

We quantified these spectral changes in 20 second, artifact-free clips selected before and after LOR and ROR using spectral edge frequency, total power, and the percentage of power at delta (1-4Hz), theta (4 – 8Hz), alpha (8 – 12Hz), beta (12-25Hz), and gamma (25 – 50Hz) frequency bands. We observed a significant decrease in spectral edge frequency from pre-to post-LOR (Figure 4A), and a significant increase in spectral edge frequency from pre-to post-ROR (Figure 4B). Additionally, we observed a significant increase in total power from pre- to post-LOR (Figure 4C), but no significant change in total power from before to after ROR (Figure 4D). This is reflected in a more uniform distribution of power across frequency bands (Figure 3), which recapitulates the emergence patterns of other GABAergic agents, like sevoflurane, reported in other studies (Chander et al., 2014). We summarize the percentage of power in band limited frequency ranges for LOR in Table 1 and for ROR in Table 2. Our pre- and post-LOR clips showed significant differences in the percentage of gamma power (Table 1). The percentage of power in the alpha range was close to significance but did not pass the correction for multiple comparisons (Table 1). Significant differences in pre- versus post-ROR clips were observed in delta (decrease), theta (decrease), alpha (decrease), beta (increase), and gamma (increase) percentages of power (Table 2). There was also an increase in beta following ROR, but it did not reach significance after being corrected for multiple comparisons (Table 2).

**Figure 4.**
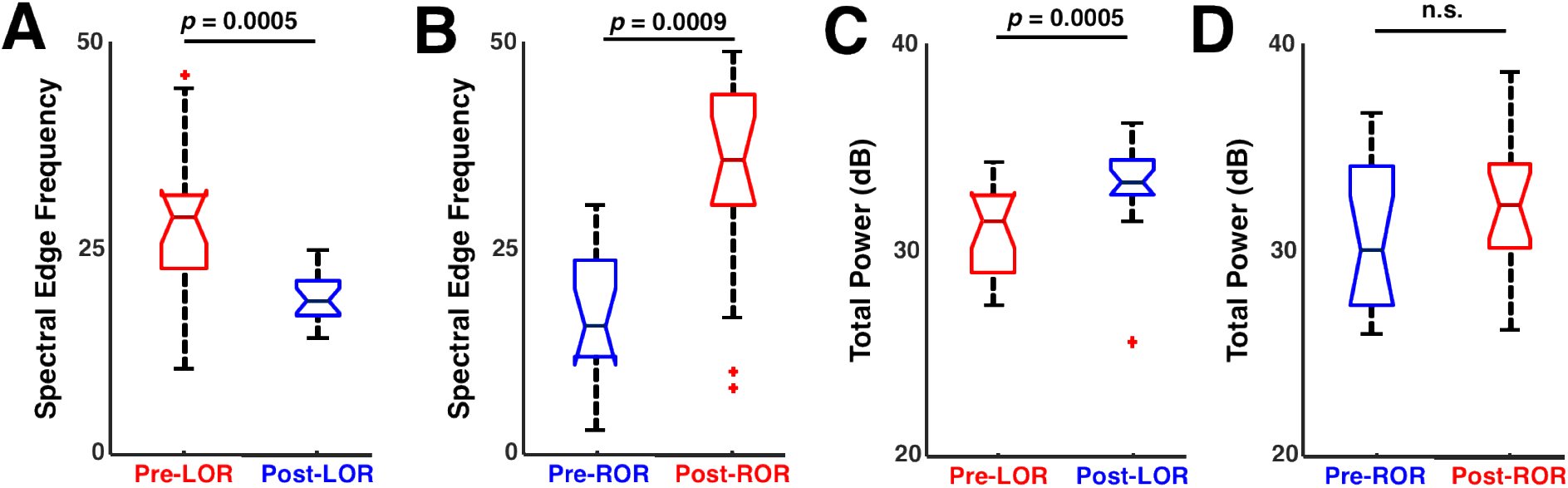
Spectral changes before and after loss and recovery of response. Spectral edge frequency and total power changes that occur in the 20 second clips analyzed before and after loss of response (LOR) and recovery of response (ROR). **A)** A significant decrease in spectral edge frequency was observed before to after LOR. **B)** A significant increase in spectral edge frequency was observed before to after ROR. **C)** A significant increase in total power was observed before to after LOR. **D)** A decrease in total power was observed before to after ROR. Reported *p* values are uncorrected.

### Attractor characterization using ellipse radius ratio and correlation dimension distinguishes between pre- and post-response transitions

To assess complexity differences before and after LOR and ROR we created three-dimensional time-delayed embeddings of the 20 second clips using a 4 ms delay (shown in Figure 1C,D, Figure 5 flattened in 2D). These embeddings appear to explore a limited amount of the possible 3D space, suggesting that they reveal attractors in the dynamics. As in previous reports, we observed flatter, more ellipsoidal attractors during periods when responses were absent and patients were anesthetized (Figure 1C,D, Figure 5). Conversely, broader, more spheroidal shapes were observed when patients were responsive (Figure 1C,D, Figure 5).

**Figure 5.**
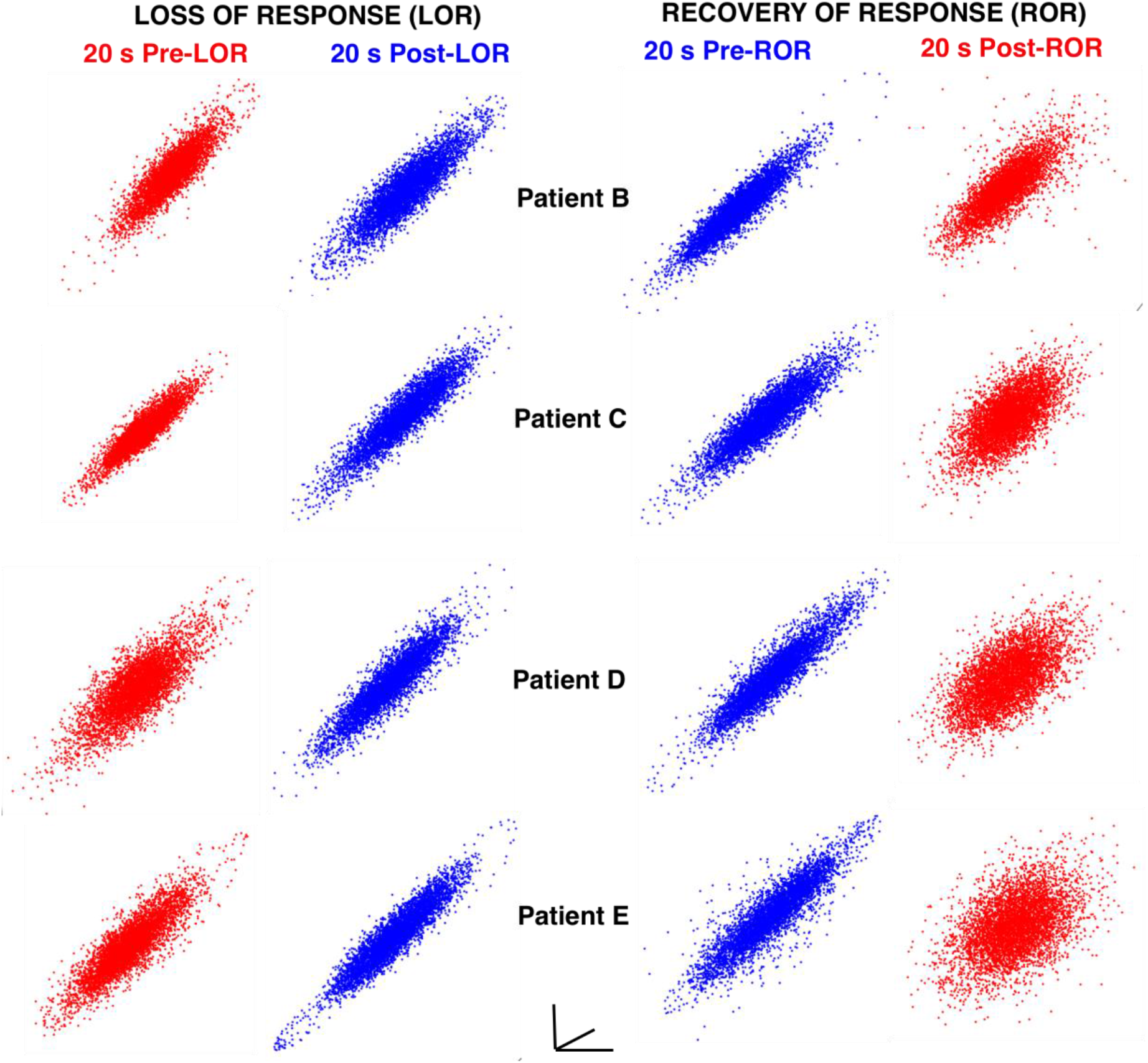
Attractor shape changes with loss and recovery of response. Examples of shape changes that occur before and after loss and recovery of response using 4 ms embedding delays. Although the attractors are three-dimensional, a two-dimensional projection is shown here for each attractor for each patient at the same viewing angle. During behavioral states when patients respond to verbal commands (red) attractors are more spherical and three-dimensional. During states when patients do not respond (blue), attractors are more ellipsoidal and flat.

To quantify the changes in the attractors observed before and after LOR and ROR, we measured the ellipse radius ratio (ERR) of the 3D attractor point clouds fit with ellipsoidal solids of revolution as previously described (Eagleman, Vaughn, et al., 2018; Eagleman, Drover, et al., 2018). Consistent with our previous work, a significant decrease in ERR was observed in the clips preceding and following LOR, while a significant increase in ERR was observed in the clips preceding and following ROR (Figure 6A). The attractor shape change was also quantified using the correlation dimension (CD). A nonsignificant decrease was observed in the correlation dimension computed using a three-dimensional embedding (3D CD) following LOR; however, a significant increase in 3D CD from before to after ROR was observed (Figure 6B). To determine if this effect occurred if the embedding dimensionality was increased, we used a five-dimensional embedding and recalculated the correlation dimension (5D CD). A significant change in 5D CD was observed in both conditions – a significant decrease pre- and post-LOR, and a significant increase pre- and post-ROR (Figure 6C).

**Figure 6.**
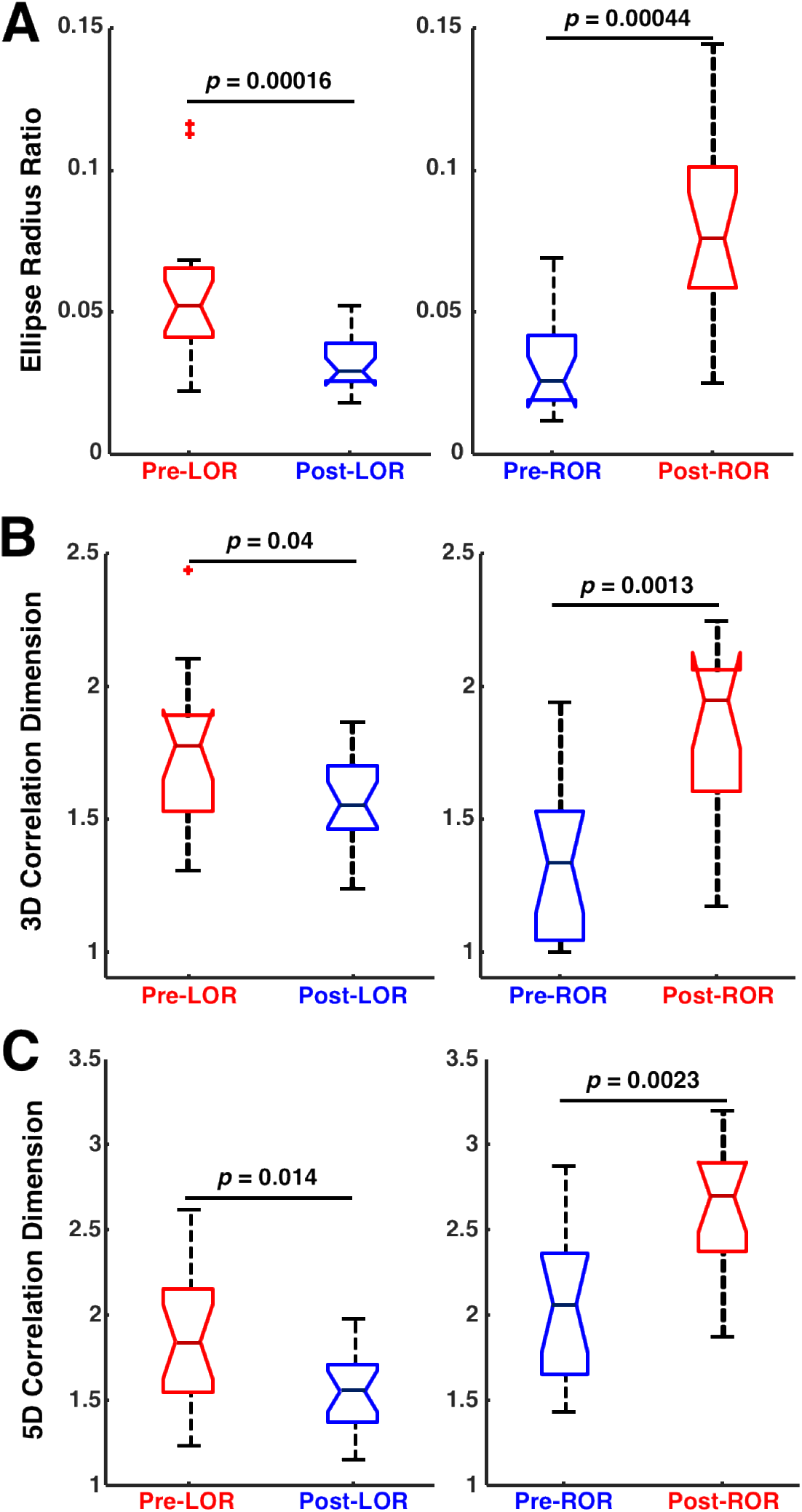
Attractor characterization measures before and after loss and recovery of response. **A)** Ellipse radius ratio (ERR) of three-dimensional attractors significantly decreases in the transition preceding loss of response (LOR) to the post-LOR state. ERR significantly increases in the transition that precedes recovery of response (ROR) to the post-ROR state. **B)** Correlation dimension (CD) of three-dimensional attractors decreases from pre-to post-LOR, but not significantly. CD increases significantly between the pre- and post-ROR state. **C)** Correlation dimension of five-dimensional attractors decreases significantly from pre-to post-LOR, and increases significantly between the pre- and post-ROR state. Reported *p*-values are uncorrected.

### Ellipse radius ratio is embedding delay dependent, but the correlation dimension is not

We explored whether these differences in ERR and correlation dimension were dependent on the embedding delay used to create the attractors. To test this, we characterized attractors using 10 successively increasing delays (4, 8, 12, 52, 100, 500, 1000, 1500, 2000, and 2500 ms, max set using autocorrelation of EEG signals). Visual inspection of the attractors shows that the obvious shape differences seen preceding and following both LOR (Figure 7A) and ROR (Figure 7B) at smaller delays become less pronounced as the delay increases, successively fading as attractors are plotted at larger delays. We assessed the significance of the measured values by delay. The ellipse radius ratio (ERR) was significant for the first three delays (*p* (corrected) = 0.0015, 0.0015, 0.011 for 4, 8, and 12 ms respectively) for loss of response, but not for the rest of the delays (Figure 8A, top). Additionally, the ellipse radius ratio (ERR) was significant for the first three delays (*p* = 0.0039, 0.0039, 0.009 corrected for 4, 8, and 12 ms respectively) for recovery of response, but not for the rest of the delays (Figure 8A, bottom). The 3D correlation dimension (3D CD) for loss of response was not significant for any of the delays (Figure 8B, top). However, for recovery of response, the 3D CD was significant for all delays (*p* (corrected) = 0.0013, 0.0045, 0.011, 0.0032, 0.0013, 0.0019, 0.0045, 0.011, 0.0013, 0.0032 for 4, 8, 12, 52, 100, 500, 1000, 1500, 2000, and 2500 ms, respectively, Figure 8B, bottom). The 5D CD was not significant for any of the delays during LOR (Figure 8C, top). Note this result is different from Figure 6C because here we are controlling for multiple comparisons, since we are now comparing 10 embedding delays. The significance value needs to be adjusted in this case. Unlike the 3D CD, the 5D CD was significant for only the first delay (*p* = 0.022 corrected for 4 ms) for recovery of response (Figure 8C, bottom).

**Figure 7.**
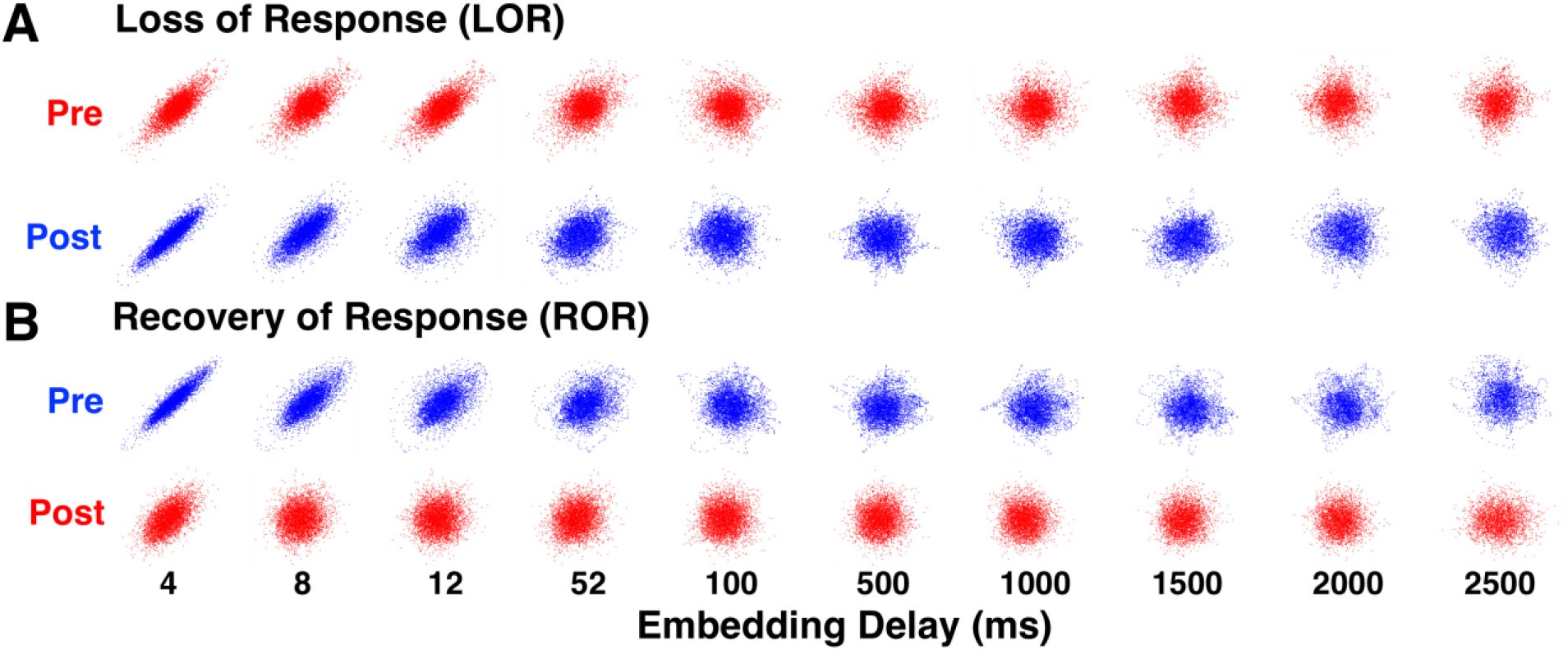
Attractors for loss and recovery of response change shape based on embedding delay. We plotted attractors from the 20 second windows we identified before and after LOR (**A**) and ROR (**B**) at a range of embedding delays. Note that the shape difference observed in the attractors disappears with increasing embedding delays. Examples are from patient D for both LOR and ROR.

**Figure 8.**
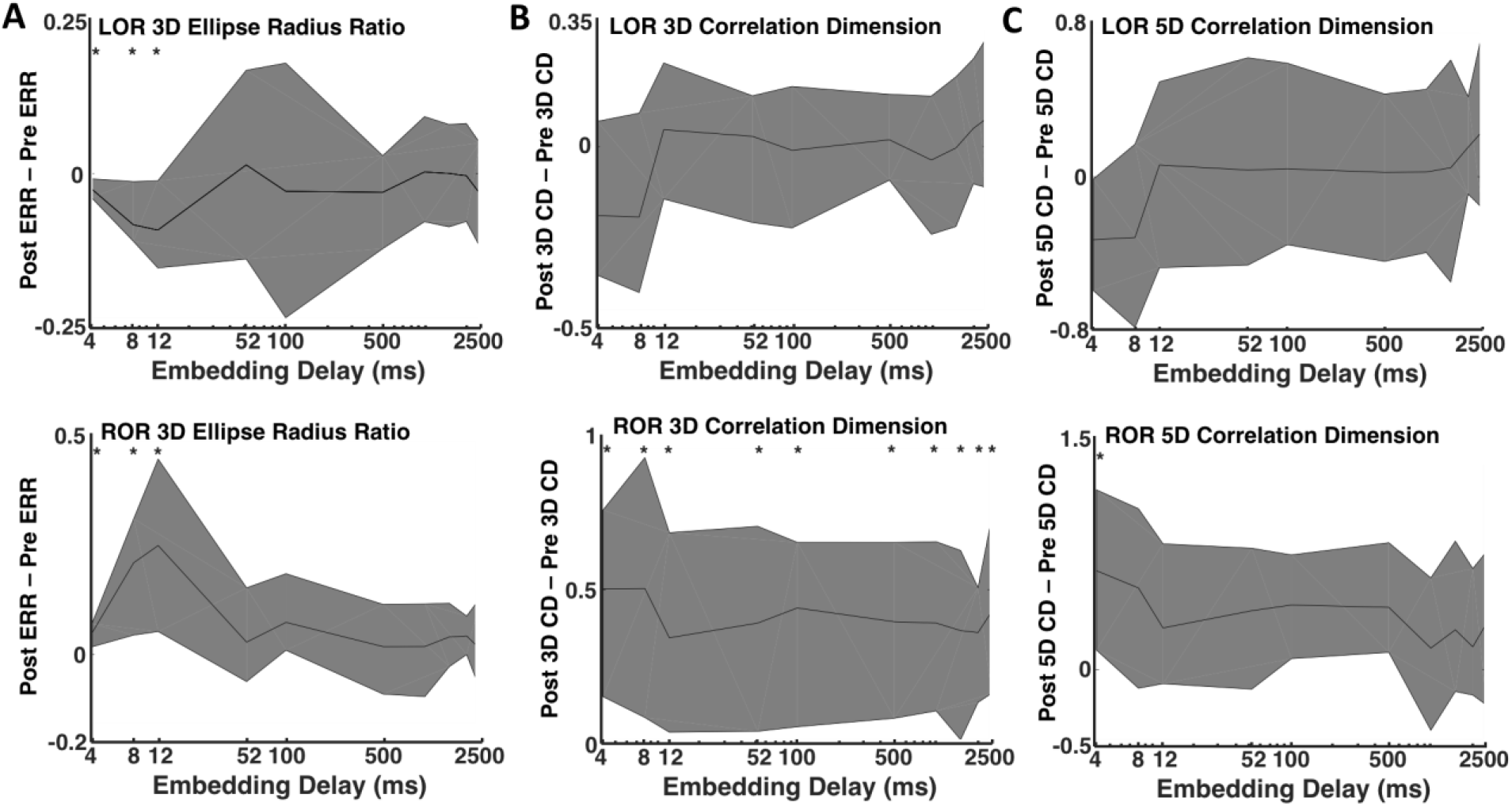
Attractor measures at increasing embedding delays for loss and recovery of response. Three-dimensional attractors were characterized using the ellipse radius ratio (ERR, **A)** and correlation dimension (CD) in both 3 **(B)** and 5 **(C)** dimensions. We calculated the differences in these three measures in 20 second EEG clips before and after LOR **(top plots)** and before and after ROR **(bottom plots)** at all embedding delays. The difference between the two clips before to after the events is shown here. Solid lines indicate the median differences for all patients at each delay and the shaded error represents the 25^th^ and 75^th^ quartiles. The * indicates a significant difference between pre- and post-clips in the EEG measures at that embedding delay at *p* < 0.05 corrected.

To reveal the scale of the dynamics of our measures and to determine whether there was a relationship between the embedding delay and the difference in our complexity measures, the Spearman correlation was calculated between the 10 embedding delays we used and the ellipse radius ratio (ERR), 3D, and 5D correlation dimensions. There were no significant correlations between the delays and ERR for LOR (r = 0.24, p = 0.51) or ROR (r = −0.66, p = 0.04). Additionally, there were no significant correlations between embedding delay and 3D correlation dimension for LOR (r = 0.62, p = 0.06) or ROR (r = −0.42, p = 0.23). The Spearman correlation was not significant (when controlling for multiple comparisons), but was strongest between the embedding delay and the 5D correlation dimension for LOR (r = 0.72, p = 0.02) and ROR (r = −0.70, p = 0.03). Given the lack of significant relationships between embedding delays and our attractor characterizations, *p*-values reported here have not been corrected for multiple comparisons.

### Attractor differences track spectral changes and patient behavioral state

To explore the dynamics of our complexity measures around our events of interest, we evaluated the two complexity measures in the 2-minute epochs preceding and following either LOR or ROR. LOR spectrograms and their corresponding attractors, calculated in 20 second adjacent windows, are shown in Figure 9. One can clearly see the attractors flattening and becoming more ellipsoidal when approaching LOR and continuing to flatten after LOR in both example patients. ERR, 3D, and 5D correlation dimensions are also shown for both patient examples in Figure 9A and B. In both individuals, the LOR trend appears to be a decline in these measures across this window, though not consistently. In these patients, ROR shows the opposite trend, with attractors inflating and becoming more spheroidal as a patient becomes responsive (Figure 10A and B). This is captured by all three attractor measures, the ERR, 3D, and 5D correlation dimensions, which all increase in the period spanning ROR. There is, however, a difference in the two patient examples shown. The patient in Figure 10A appears to be moving to a more unresponsive state at approximately minute +1.75, indicated by the decrease in higher frequencies and reappearance of a slow-wave dominated spectrogram. The attractors follow this trend during this period and become more ellipsoidal and flat. In the patient in Figure 10B, however, we do not see this reversion, reflecting a behavioral state in which the subject remained responsive. When we explore the dynamics of these measures for all patients, we can see a generally decreasing trend across the LOR time window and an increasing trend across the ROR time window (Figure 11). The LOR dynamics appear to change more gradually and consistently, while the ROR appears to have an abrupt jump in the complexity measures once the patient regains their ability to respond (Figure 11).

**Figure 9.**
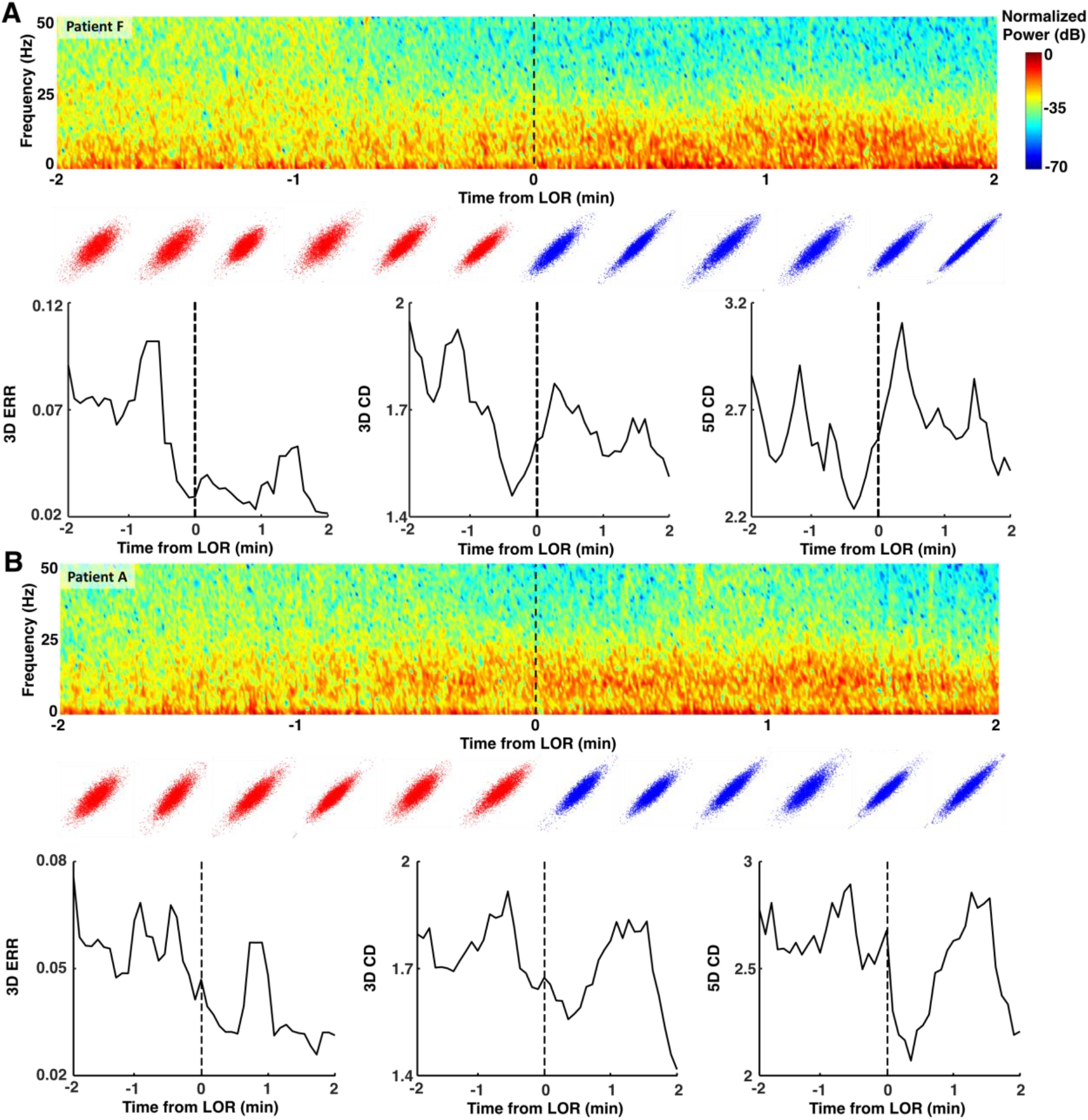
Examples of dynamics of attractors around loss of response (LOR). Spectrograms and attractors for epochs surrounding LOR by +/− two minutes for two individual patients (**A and B**). Attractors are three-dimensional but shown projected into two dimensions from the same viewing angle. Note the flattening and more ellipsoidal shapes approaching and following LOR in both patients. The ellipse radius ratio (ERR) measure appears to be the only one to decrease throughout the LOR period.

**Figure 10.**
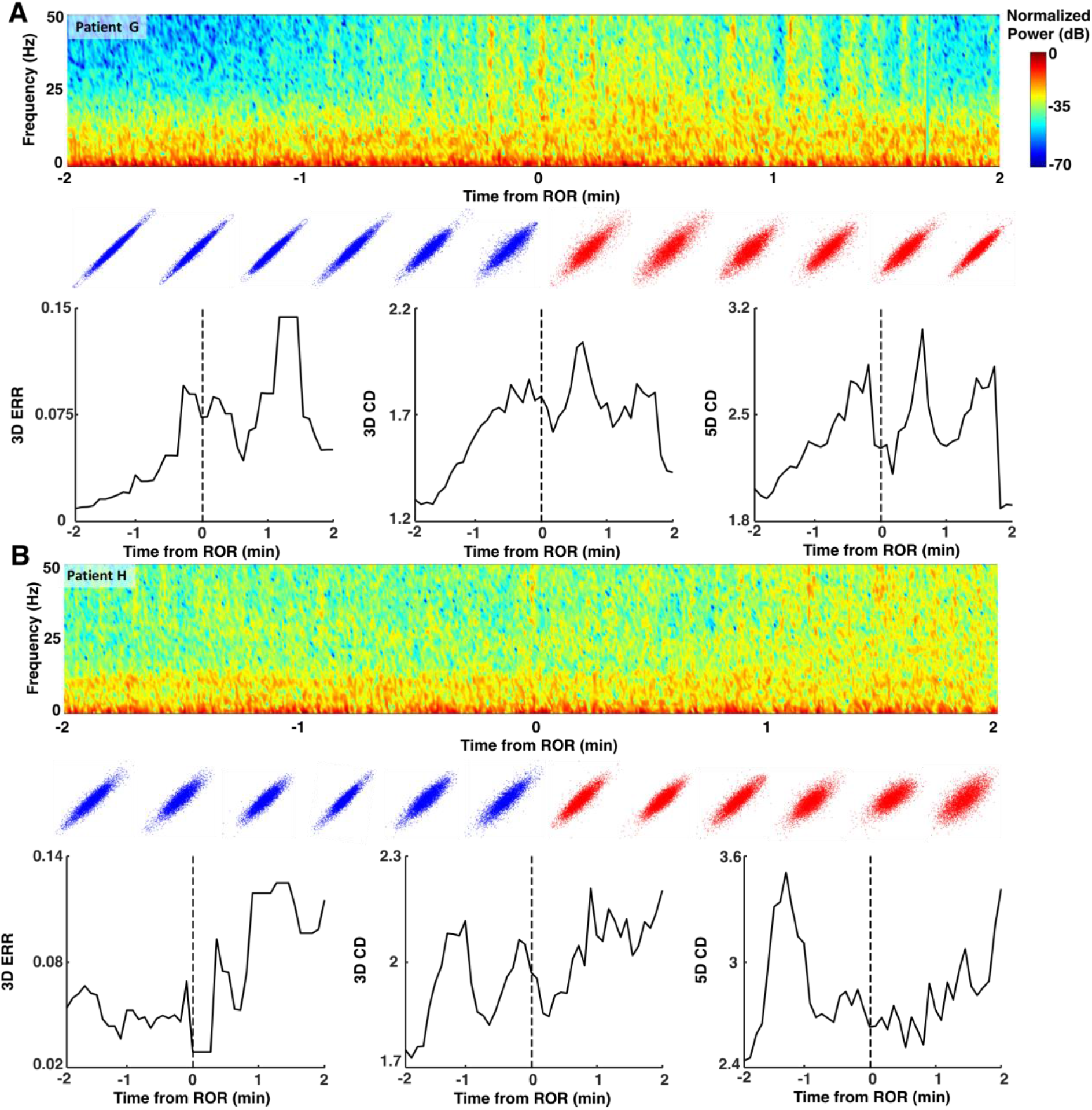
Examples of dynamics of attractors around recovery of response. Spectrograms and attractors for epochs surrounding ROR by +/− two minutes for two individual patients (**A and B**). Attractors are three-dimensional but shown projected into two dimensions from the same viewing angle. Note the broadening and more spheroidal shapes approaching and following ROR in both patients. In patient G (**A**), at approximately +1.75 minutes, the ellipsoidal attractor begins to flatten again, reflecting the patient’s behavioral state moving to becoming less responsive again with lack of stimulation.

**Figure 11.**
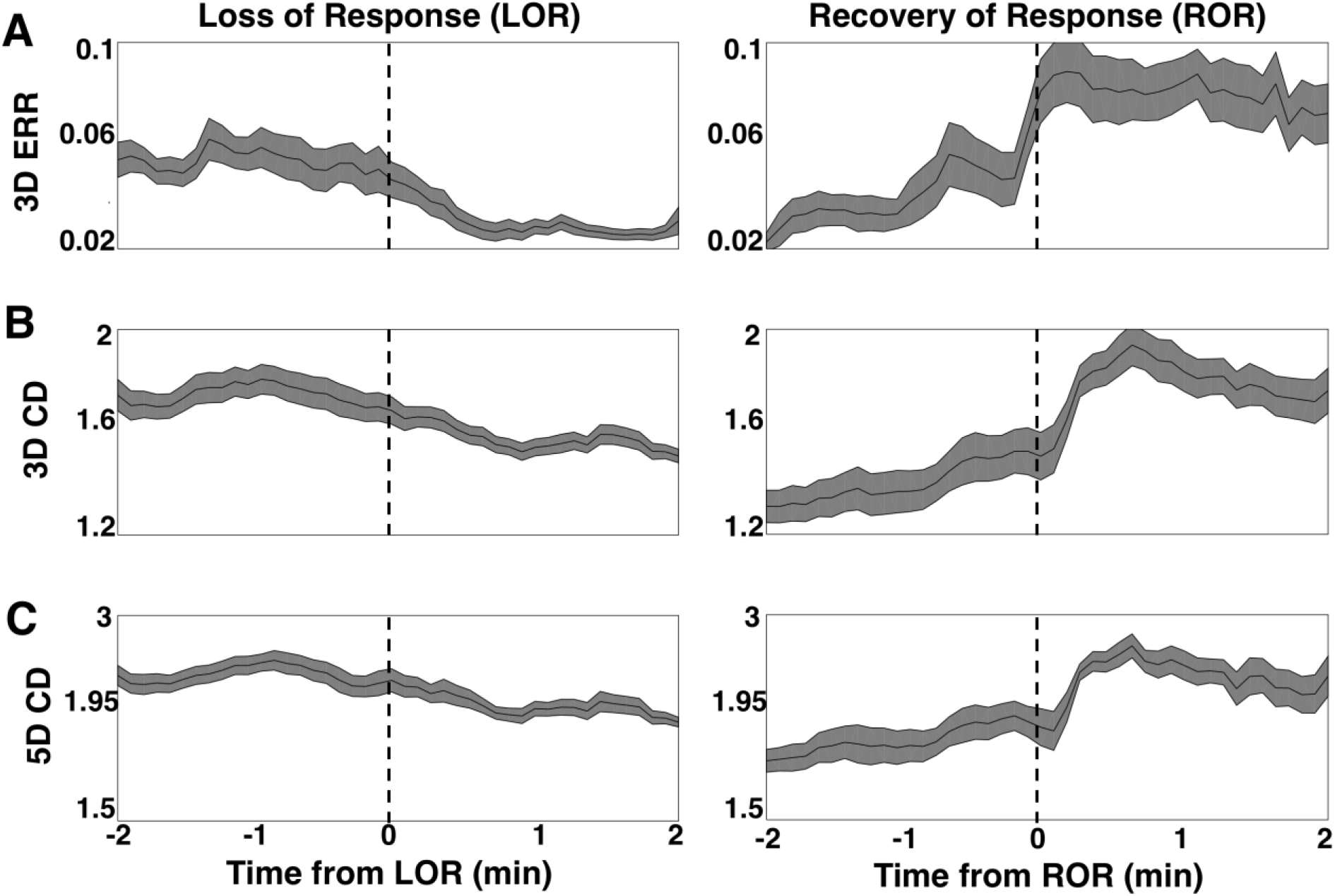
Dynamics of attractors during loss and recovery of response for the population studied. **A)** The average ellipse radius ratio (ERR) for all patients appears to decrease gradually as patients approach loss of response (LOR), and continues to decrease in value as anesthetic depth increases **(A, left)**. In contrast ERR upon recovery **(A, right)** is marked by a more abrupt increase in ERR at the ROR transition point. **B)** The 3D correlation dimension (CD) gradually decreases during LOR **(B, left)** and more abruptly increases during ROR **(B, right). C)** The 5D correlation dimension (CD) gradually decreases during LOR **(C, left)** and more abruptly increases during ROR **(C, right)**. Both the 3D CD ROR **(B, right)** and the 5D CD ROR **(C, right)** appear to lag behind the recovery timepoint (dashed line), while the ERR does not. Solid black lines represent the mean complexity values at each timepoint for all patients. Shaded area is the standard error of the mean. Vertical dashed lines represent the time of LOR or ROR.

### Complexity measure effect sizes are comparable to spectral measures and strongly correlated with gamma

To determine the effect size of our EEG measures we calculated pairwise Cohen’s D for spectral edge frequency, total power, band-limited frequency ranges, and complexity measures (Table 3). This measure gives a description of the difference of means, with +/−1 reflecting a large effect size. The ERR phase-space analysis, spectral edge frequency, total power, and gamma power reflected the largest effect sizes for LOR, of approximately 1 (Table 3, middle column). Alpha power, 5D correlation dimension, and 3D correlation dimension had medium effect sizes of 0.6-0.7. For ROR the largest effects sizes were ERR (1.4) and spectral edge frequency (1.3), followed closely by 3D (1.2) and 5D (1.0) correlation dimension estimates (Table 3, last column). While these effect sizes for ROR were greater than LOR, the total power effect size was much smaller in the ROR condition (Table 3). Effect sizes of changes in power in all the frequency bands were relatively larger for ROR, with the exception of the gamma band (Table 3). To determine whether there was a relationship between our EEG measures we calculated the Spearman correlation. Specifically, the Spearman correlation between the pre- and post-event complexity measures (ERR, 3D CD at 4 ms delay) and percentage of power differences in individual frequency bands was computed separately for LOR (Table 4) and ROR (Table 5). During LOR, there was a significant relationship between the change in ERR and gamma, as well as between 3D CD and gamma (Table 4). Similarly, during recovery of response (ROR), significant correlations were observed between ERR and gamma as well as 3D CD and gamma (Table 5). Interestingly, the correlation between ERR and 3D CD reached significance for the ROR condition, even when corrected for multiple comparisons; it did not rise to significance for the LOR condition.

**Table 3.** Summary of Cohen’s D for measured values before and after loss of response and recovery of response. Cohen’s D (sign convention: post – pre) was calculated to quantify the effect sizes of various EEG measures by comparing means. We included all measures that showed significant differences before and after LOR or ROR, and the standard frequency bands for comparison. ERR, and CD values (for both dimensions) are shown here for 4 ms (shortest) delay.

| EEG measure | Cohen's D LOR | Cohen's D ROR |
| --- | --- | --- |
| SEF | -1.1 | 1.3 |
| Total Power | 1.0 | 0.38 |
| Delta | -0.18 | -0.80 |
| Theta | -0.12 | -0.78 |
| Alpha | 0.68 | -0.79 |
| Beta | 0.23 | 0.58 |
| Gamma | -0.95 | 0.87 |
| ERR | -1.1 | 1.4 |
| CD 3D | -0.59 | 1.2 |
| CD 5D | -0.67 | 1.0 |

**Table 4.** Summary of correlations for measured changes before and after loss of response (LOR). Spearman Correlation calculated between pairs of EEG measures. ERR and CD 3D, calculated for a 4 ms delay, were compared against each other as well as percentage of power in each of the frequency bands. The * indicates that the correlation is significant at *p* < 0.05 corrected.

| Parameter 1 | Parameter 2 | Spearman's Rho | Significance value, corrected |
| --- | --- | --- | --- |
| ERR | CD (3D) | 0.54 | 0.07 |
| ERR | Delta | -0.12 | 0.79 |
| ERR | Theta | -0.23 | 0.61 |
| ERR | Alpha | 0.019 | 0.94 |
| ERR | Beta | 0.16 | 0.79 |
| ERR | Gamma | 0.71 | 0.0055* |
| CD (3D) | Delta | -0.28 | 0.53 |
| CD (3D) | Theta | -0.48 | 0.11 |
| CD (3D) | Alpha | 0.054 | 0.91 |
| CD (3D) | Beta | 0.11 | 0.79 |
| CD (3D) | Gamma | 0.73 | 0.0055* |

**Table 5.** Summary of correlations for measured changes before and after recovery of response (ROR). Spearman Correlation calculated between pairs of EEG measures. ERR and CD 3D, calculated for a 4 ms delay, were compared against each other as well as percentage of power in each of the frequency bands. The * indicates that the correlation is significant at *p* < 0.05 corrected.

| Parameter 1 | Parameter 2 | Spearman's Rho | Significance value, corrected |
| --- | --- | --- | --- |
| ERR | CD (3D) | 0.70 | 0.012* |
| ERR | Delta | -0.41 | 0.12 |
| ERR | Theta | -0.54 | 0.075 |
| ERR | Alpha | -0.45 | 0.11 |
| ERR | Beta | 0.51 | 0.084 |
| ERR | Gamma | 0.91 | 0* |
| CD (3D) | Delta | -0.48 | 0.10 |
| CD (3D) | Theta | -0.41 | 0.12 |
| CD (3D) | Alpha | -0.35 | 0.19 |
| CD (3D) | Beta | 0.59 | 0.0495* |
| CD (3D) | Gamma | 0.73 | 0.0099* |

## DISCUSSION

Our results agree with previous reports of the utility of complexity analyses to discriminate between the subtle EEG changes that occur in operating room patients during induction (Eagleman, Vaughn, et al., 2018; Eagleman, Drover, et al., 2018; Watt & Hameroff, 1988) and emergence (Eagleman, Drover, et al., 2018; Walling & Hicks, 2006) from anesthesia. We demonstrate here that these approaches provide equivalent discrimination between periods preceding and following LOR, and improved discrimination between periods preceding and following ROR. As in our previous work, our geometric phase-space ellipse radius ratio (ERR) was significantly different between pre and post-LOR and ROR clips. Additionally, significant differences were found using 3D correlation dimension between pre and post-ROR. We also found that our ERR values were correlated with correlation dimension values during ROR. Since ERR is correlated with multiscale entropy (Eagleman, Vaughn, et al., 2018), another measure of complexity, and the correlation dimension used here, we reason that ERR may itself capture signal complexity. In fully awake patients, the attractor is a nearly perfect sphere. As patients lose consciousness, the attractor collapses into an ellipsoidal shape and explores less of the possible trajectory shape, thus decreasing complexity and information content. After LOR a clear additional compression of the attractor was evident when comparing the 20-second moving windows of EEG before and after patients stopped responding to verbal command. The opposite trend is observed upon ROR indicating a recovery of complexity and information upon arousal. This finding confirms a previous report of correlation dimension increase with ROR from anesthetic administration (Walling & Hicks, 2006). Additionally, we also observed an abrupt change in the correlation dimension (as well as our other measures) near the ROR transition as has been previously described (Walling & Hicks, 2006).

In this data set, attractors following ROR were observed to be more spheroidal than prior to LOR (Figure 5). This is a function of the propofol infusions being started significantly before (i.e., minutes) we started calculating the attractors in the LOR condition. We wanted to compare close time windows to significant anesthetic depth transitions (loss of consciousness) to test our measures on subtle EEG changes. The difference in pre-LOR and post-ROR attractor shape, along with the attractor shape changes that occur over LOR (Figure 9) and ROR (Figure 10), shows that the ellipse radius ratio measure closely follows the dynamics during these events. This is especially true in Patient G (Figure 10A), where the patient becomes less responsive again with lack of stimulation after initial recovery of consciousness. This relapse to unconsciousness is not uncommon in patients who have emerged from anesthesia but are no longer being stimulated. It is as if their states of wakefulness and consciousness oscillate. Also, many drugs such as propofol and the more lipophilic volatile anesthetics have a context-sensitive half-life (Struys et al., 2000) and remain in fatty tissue. They form a depot or reservoir of anesthetic that continues to leach out even after the maintenance anesthetic is terminated. The careful tracking of our method during this period of recovery means these methods could be useful to assess level of consciousness in the recovery post-anesthesia care unit as an added measure of vigilance and safety.

Unlike previous reports (Eagleman, Vaughn, et al., 2018; Eagleman, Drover, et al., 2018) where complexity measures show enhanced performance over spectral measures, here our complexity measures had equivalent performance compared to spectral measures. We reason that this is due to the use of propofol with adjuvant agents (e.g. remifentanil, fentanyl, midazolam) which enhances the signal change from high frequency, low amplitude awake activity to low frequency, high amplitude activity upon anesthetic induction (Billard et al., 1997; Egan, 1995; Greenblatt et al., 2004; Minto et al., 1997; Scott, Ponganis, & Stanski, 1985). In contrast, use of agents like nitrous oxide with propofol make spectral measurements less reliable (Foster & Liley, 2011; Hirota, Kubota, Ishihara, & Matsuki, 1999; Rampil, Kim, Lenhardt, Negishi, & Sessler, 1998). Here the predominant anesthetic was a slow infusion of propofol; thus, spectral measures were expected to discriminate between the two states well.

Previous reports indicate significant differences in alpha power following induction of anesthesia (Akeju et al., 2014; Feshchenko, Veselis, & Reinsel, 2004; P. L. Purdon et al., 2013). We found significant differences in geriatric patients induced with propofol and fentanyl (Eagleman, Vaughn, et al., 2018). However, here we found an increase but not significant difference in the percentage of alpha before and after LOR (Table 1), but did observe a significant reduction upon ROR (Table 2). Our time points are very close to the clinically relevant loss of response and propofol anesthesia was delivered slowly and gradually over several minutes, whereas in our previous study (Eagleman, Vaughn, et al., 2018) propofol was delivered in a bolus so LOR and EEG changes were likely achieved faster than those observed here. Thus, the dynamic state of the brain around LOR was not so greatly altered during the time points investigated.

Since we are retrospectively analyzing previously collected data, we have some limitations in interpretation. We restricted our analysis to patients who were maintained on propofol infusions; however, adjuvant agents that could influence the EEG, including lidocaine, remifentanil, midazolam, and fentanyl, were also co-administered. In humans, slowing of EEG activity from high frequency, low amplitude signals to low frequency, high amplitude signals is observed when administering these agents (Billard et al., 1997; Egan et al., 1993, 1996; Mi, Sakai, Kudo, Kudo, & Matsuki, 2003; Veselis, Reinsel, Marino, Sommer, & Carlon, 1993). This is reflected by significant observations in increased total power (Kortelainen, Koskinen, Mustola, & Seppänen, 2009), delta power (Kortelainen et al., 2009), and decreased spectral edge frequency (Egan et al., 1996). When combined with propofol for sedation, remifentanil decreases the amount of anesthetic needed and can allow for lighter sedation levels while maintaining patient comfort (Mazanikov et al., 2011). Additionally, a similar decrease in SEF was observed after patients lost response to verbal commands (LOR); however, addition of remifentanil (compared to propofol only) attenuated this effect in a dose-dependent manner (Kortelainen, Koskinen, Mustola, & Seppänen, 2008). It also appears to attenuate the increase in delta activity following LOR compared to the administration of propofol alone (Kortelainen et al., 2009). Lidocaine decreases the median frequency when added to propofol anesthesia (Kaka et al., 2015; Maher, Reynolds, & Chander, 2016). Co-administration of fentanyl with propofol anesthesia decreases the amount of propofol needed to keep patients unconscious but does not influence spectral measures upon emergence (Mi et al., 2003). All of these agents appear to influence the EEG dynamics in the same manner as propofol relatively, driving the EEG to slower, higher amplitude patterns. Additionally, the anesthesia used in patients in this study represents clinically relevant administration protocols. The ability of these complexity measures to follow the consciousness transitions during routine anesthesia care highlights their utility.

In summary, we have demonstrated the utility of complexity measures to distinguish between the subtle EEG changes that occur with loss and recovery of response. On a larger scale, our work fits in with other studies from the neuroscience community showing that complexity changes occur in neural activity with consciousness transitions (Bodart et al., 2017; Casali et al., 2013; Rosanova et al., 2018; Sarasso et al., 2015). As the brain naturally transitions from wakefulness to sleep or has unconsciousness imposed with small molecules as in anesthesia, neural activity becomes more synchronous. Decreases in functional connectivity with anesthesia (Lee, Mashour, Noh, Kim, & Lee, 2013; George A. Mashour & Hudetz, 2018) impose regularity, periodicity, and consistency. Complexity measures of EEG signals capture these changes well, especially in patients who exhibit behavior associated with unconsciousness. These observations in our work, consistent with others, highlight that complexity measures from nonlinear dynamics not only have utility as measures of anesthetic depth, but also support existing theories on neural correlates of consciousness transitions in the brain.

## MATERIALS AND METHODS

### Patient Selection

A retrospective analysis of EEG data as part of a medical chart review was approved by the Stanford University School of Medicine Institutional Review Board Protocol 28130. Twenty patients were chosen from a population of surgical all-comers to the operating room at Stanford Hospital and Clinics. Patients were included if they were to be anesthetized with propofol as their maintenance anesthetic (a total intravenous anesthetic or TIVA). Subjects were between the ages of 23 and 87 years old (median age 51), comprised American Society of Anesthesiology (ASA) classes 1-3, and were undergoing various surgical procedures (breast, gynecological, head and neck, and neurosurgery). Subjects included 17 females and 3 males. We deliberately did not restrict the population studied to a single surgical cohort, in order to better reflect the diversity of patients that would undergo surgical anesthesia, and to demonstrate that nonlinear dynamics tools would be sensitive to brain state.

### Anesthetic Administration

Patients were administered an infusion of intravenous propofol over minutes in order to achieve a slowly increasing plasma concentration of anesthetic for hemodynamic stability on induction of anesthesia, and to more accurately determine the precise time of the loss of response (LOR). LOR was defined at the moment the patient lost response to verbal stimulus (eye opening or verbal response to name), assessed every 5 seconds after the infusion was started. If there was doubt regarding the LOR, an additional verbal stimulus was delivered to the patient. Recovery of response (ROR) was similarly assessed by verbal stimulus (stating the patient’s name and asking them to open their eyes) after the maintenance anesthetic was turned off. Timing of LOR and ROR were marked to the nearest second for off-line analysis of EEG signals. We estimate an error range for the observer to be less than 5 seconds based on continuous vigilance of the patient between stimuli. Raw EEG data, processed EEG data (patient state index or PSI), and time stamps were downloaded for off-line data analysis.

Choice of airway device (laryngeal mask airway or endotracheal tube) for general anesthesia was determined by the anesthesiologist based on the type of surgical procedure and patient needs. LOR measurement was unaffected since it occurred prior to instrumentation of the airway. Patients were maintained on propofol infusions as the maintenance anesthetic (TIVA), with analgesic adjuncts as needed according to the anesthesiologist’s clinical judgment. Some patients received pre-operative medication (midazolam and/or fentanyl) prior to initiation of the propofol infusion as clinically necessary. Intraoperative analgesia was supplied by fentanyl, hydromorphone, ketorolac, acetaminophen, or ketamine IV boluses, and supplemented by lidocaine, fentanyl, or remifentanil infusions according to the anesthesiologist’s discretion and the degree of surgical stimulation. Timing of ROR would have been influenced by maintenance anesthesia and analgesic adjuncts, but was equally compared across all analyses.

### EEG Data Acquisition and Preprocessing

Standard American Society of Anesthesiologists (ASA) monitors (pulse oximeter, non-invasive blood pressure cuff, capnograph, electrocardiogram) (Committee, 2015) were utilized for each patient in order to measure intraoperative oxygen saturation, hemodynamic parameters, and end-tidal gas concentrations (O2, CO2). A self-adhesive, 5-lead frontal EEG electrode array was placed according to the manufacturer’s instructions (SedLine Legacy, Masimo, Irvine, CA) on the forehead of each patient prior to induction of anesthesia. Electrodes were applied at approximately F7, F8, Fp1, and Fp2 referenced to AFz (Figure 1A) in the standard 10-20 electrode EEG montage. Appropriate electrode impedance was checked according to an automated SedLine routine that sends a ~78 Hz impedance pulse to each electrode. Data was digitized at 250 Hz. EEG recordings from F7 were de-trended and then filtered using a 50 Hz low-pass Butterworth Infinite Impulse Response (IIR) filter, and a 1 Hz high-pass Butterworth IIR filter for subsequent analysis.

Identification of clips for analysis has been previously described (Eagleman, Vaughn, et al., 2018; Eagleman, Drover, et al., 2018). Briefly, 20 second artifact-free clips occurring within 2 minutes before to 2 min after loss and recovery of response to verbal commands (LOR and ROR respectively) were selected manually for subsequent analysis (Figure 1A). Two of the authors visually inspected the EEG traces as well as the EEG spectrograms to ensure the clips did not contain burst suppression or artifacts (SLE, MBM). From the original 20 patients, 19 were selected for LOR analysis and 16 selected for ROR analysis given these were the patients who had artifact-free EEGs for at least 20 continuous seconds before and after the events of interest.

### Spectral Analysis

EEG Data was analyzed using custom scripts written in MATLAB (Mathworks, Natick, Massachusetts). Power spectral density was calculated on the 20 second EEG clips during the pre- and post-LOR and pre- and post-ROR periods using the multitaper method as part of the ‘mtspectrumc’ function of the Chronux toolbox (Mitra & Bokil, 2008) (http://chronux.org/). Specifically, we used a time-bandwidth product of 5, 9 tapers, and limited the frequency range to 0 to 50 Hz. The error range for power spectral density plots was computed using the theoretical estimate method at 0.05 significance level. Power values were expressed in decibels.

To visualize the temporal profiles of the spectral changes occurring during the 4 minute windows surrounding the LOR and ROR transitions (i.e. 2 minutes before to 2 minutes after), we computed normalized spectrograms with custom MATLAB scripts (Figure 1B). A Fourier transform (using ‘fft’ function in MATLAB) was performed using Hann windows with half window overlap. The magnitude was converted to decibels (dB) and the spectrogram was normalized by its maximum magnitude.

Total power and spectral edge frequency (SEF, the frequency below which 95% of the total EEG power resides) were also calculated using multitaper spectral analysis (Mitra & Bokil, 2008) on the filtered EEG. In addition, we calculated the percentage of total power for individual frequency bands per condition (pre- and post-LOR, pre- and post-ROR). The percentage of total power was used because significant changes in EEG total power with exposure to propofol have been previously reported. The frequency band ranges were defined as follows: delta: 1 to 4 Hz; theta: 4 to 8 Hz; alpha: 8 to 12 Hz; beta: 12 to 25 Hz; and gamma: 25 to 50 Hz (Gugino et al., 2001; P. L. Purdon et al., 2013).

### Characterization of Dynamical Attractors

#### Geometric Phase-Space Analysis

We constructed three-dimensional, time-delayed embeddings (attractors) of the EEG signal before and after LOR and ROR using a 4 ms delay for correlation dimension analysis (Figure 1C). We chose this delay as it was the smallest delay possible for our sampling frequency and showed the most significant effects based on previous work (Eagleman, Vaughn, et al., 2018; Eagleman, Drover, et al., 2018). We observed shape changes in attractors before and after our time points of interest when plotted at this timescale. We quantified this shape change by fitting the three-dimensional attractor to an ellipsoidal solid of revolution (Eagleman, Vaughn, et al., 2018; Eagleman, Drover, et al., 2018; Khachiyan, 1980) (Figure 1D). The lengths of the symmetry axes of the ellipsoid were calculated and the ratio of the minimum and maximum axes (which we term the ellipsoid radius ratio, ERR) was used to quantify the shape change. A radius ratio of 1 implies a sphere, while smaller ratios imply more strongly ellipsoidal shapes.

#### Correlation Dimension

Using the same three-dimensional attractor with 4 ms embedding delay (Figure 1C), we tested whether the correlation dimension captured the EEG changes that occurred before and after LOR and ROR. We used a previously reported and commonly used algorithm to compute the non-integer (fractal) correlation dimension that works for irregularly sampled objects (e.g., a point cloud in our case) (Grassberger & Procaccia, 1983; Walling & Hicks, 2006; Widman, Schreiber, Rehberg, Hoeft, & Elger, 2000). We also tested whether significant differences in correlation dimension could be observed when we increased the dimensionality of the embeddings from 3 to 5 dimensions.

#### Complexity Measures by Embedding Delay

We tested also whether the ellipse radius ratio and correlation dimension were changed by varying the embedding delay time. To estimate the optimal delay for creating the attractors we calculated the first zero-crossing of the autocorrelations of the pre-LOR and post-LOR signals. We used this value to set our maximum range, and tested multiple embedding delays (4, 8, 12, 52, 100, 500, 1000, 1500, 2000, and 2500 ms) between the shortest time window (shifting the EEG by 1 point) to the largest (set by the autocorrelation zero crossing). Spearman’s correlation was used to determine whether a trend existed between the embedding delays and ellipse radius ratio or correlation dimension. We tested whether a trend existed for both 3D and 5D time-delayed embeddings and correlation dimension.

#### Dynamics of Complexity Measures

To explore the dynamics of our measures around our two clinically relevant time periods (LOR and ROR) we calculated ERR and correlation dimension in a 4 min window surrounding these events. Specifically, 20 second windows of EEG activity starting 2 min before an event to 2 min after the event were used to create attractors. We shifted these 20 second windows every 5 seconds so windows had 75% overlap. We created the attractors using the smallest embedding delay (4 ms) as this delay showed the most significant differences before and after LOR and ROR. We then calculated the ERR and correlation dimension (both 3D and 5D) for each 20 second segment of EEG activity. The average and standard error of the mean of all EEG data from 19 LOR and 16 ROR patients were calculated to show the dynamics of our measures across this period.

### Effect Size and Correlations of EEG Measures

We calculated a paired-data Cohen’s D on our EEG measures before and after LOR and before and after ROR for the complexity (ERR and correlation dimension) measures. We also calculated the Cohen’s D for the spectral edge frequency, total power, and percentage of power in each of the frequency bands (delta, theta, alpha, beta, and gamma) for comparison. We used the 4 ms embedding delay complexity measures as these showed the most significant results. Additionally we calculated Spearman correlations between the ERR, 3D, and 5D correlation dimension measures and age to determine if these patient characteristics influenced our results. We also calculated the Spearman correlation between the ERR and 3D correlation dimension (again at the shortest 4 ms delay) with each other and with the percentage of power in the band-limited frequency ranges. We calculated these correlations independently for LOR and ROR periods.

Since we had a large age range in our study, we tested whether patient age may have influenced our measures. To determine whether complexity measures change with age we calculated the Spearman correlation between age and change in pre-to post-ERR and 3D CD (at 4 ms delay) measures individually for LOR and ROR. There were no significant correlations in the change in ERR or 3D CD and age for LOR (ERR and age r = 0.17, *p* = 0.48; 3D CD and age r = −0.21, *p* = 0.38, *p* values are uncorrected) and ROR (ERR and age r = 0.006, *p* = 0.98; 3D CD and age r = 0.04, *p* = 0.88, p values are uncorrected). Given the lack of correlation in these measures we reasoned that our complexity EEG measures were not confounded by age.

### Statistics

Significance values for multiple comparisons within a particular analysis type were corrected using the Holm-Bonferroni method—a sequentially-rejective procedure(Holm, 1979). Specifically, we corrected *p* values for pre-vs post-metrics for LOR and ROR separately within each of the following analyses: (1) the power percentage across all 5 frequency bands (delta, theta, alpha, beta, gamma), (2) the ellipse radius ratio (ERR) when comparing across all 10 embedding delays, (3) the correlation dimension when comparing across all 10 embedding delays, and (4) the Spearman correlation of ERR, correlation dimension, and frequency band power. We report our results as medians (25, 75 percentiles), and corrected significance values (*p*) are calculated from Wilcoxon Signed Rank Tests.

## ACKNOWLEDGEMENTS

We thank Dr. David Drover for helpful discussions with our data analysis and on identifying artifact-free EEG segments using the electroencephalogram changes that occur with propofol anesthesia administration.

## Author Contributions

MBM and SLE conceived of the study. DC collected the data. DC, SLE, and CD organized the data for analysis. SLE, MBM, and NTO analyzed the data. SLE, DC, and MBM wrote the manuscript. All authors contributed intellectually with rounds of review. All authors approved the final manuscript for publication.

## Ethics

Data was collected, de-identified, and analyzed as part of a retrospective chart review under approved Stanford University Institutional Review Board Protocol 28130.

